## Supplementary materials for "Physiological Synchrony Predicts Observational Threat Learning in Humans"

#### Supplementary methods

##### Participants

To determine sample size we simulated data. We targeted an effect size of 0.04 (in $\sqrt{\mu S}$) for the target interaction between a CRQA metric and CS status, with a standard deviation of 0.015 for the per participant varying coefficients. Model priors were same as for our subsequent analyses. We analyzed 400 simulated datasets and assessed if $BF_{10} > 3$. Our simulations indicated that 65 dyads would provide 90% power to assess an effect.

##### Equipment

Electrodermal activity was recorded using a BIOPAC MP150 system and AcqKnowledge software as a skin conductance signal measured in microsiemens ($\mu S$). Recording was at 1000Hz from the distal phalanges of the middle and index fingers from the hand belonging to the arm not receiving shocks. Ag-AgCl electrodes were used together with a conducting gel, following standard recommendations [1]. Electric shocks, consisting of a single 100ms DC pulse, were administered using a Biopac STM200 module (Biopac Systems Inc.) applied to the lower forearm. The strength of the electric shocks was individually calibrated so that participants experienced the shocks as being "unpleasant but not painful". Stimulus presentation and shock administration was controlled by scripts programmed in PsychoPy [2].

*

#### Signal processing

Electrodermal activity was continuously measured from both participants throughout the experiment. All processing of the electrodermal recordings was done in AcqKnowledge 4.1 (Biopac Systems Inc.). The raw signal from each participant was filtered offline in AckKnowledge with a low-pass filter (1Hz) to remove potential recording artefacts and then a high-pass filter (0.05Hz) to recover the phasic skin conductance responses by removing the tonic component of the signal [3]. Using CS onset and shock delivery as event markers and following established protocols [4], skin conductance responses (SCRs) were measured as the largest peak-to-peak amplitude difference in the phasic skin conductance signal in the 0.5 to 4.5 second window following stimulus onset. Responses below $0.02\mu S$ were scored as zero. Scoring was first done using AcqKnowledge's automated scoring algorithm, and then manually checked by an experimenter. SCRs were square-root transformed prior to analysis [5].

#### Procedure

Participants first read and signed consent forms. They were then given separate written instructions about their initial role during the experiment; one participant read instructions pertaining to the role of "observer" and the other participant read instructions to role of "demonstrator". Both participants were taken to the recording room and sat on chairs positioned so that they would afford a good view both of a computer screen and the other participant (see Supplementary Fig. 1). Both participants were asked to place their respective instruction sheets in front of them as a reminder of their current role. Each participant's instructions also emphasized that all communication between them was forbidden for the duration of the experiment.

The experiment was divided into four blocks and followed established protocols for video-based observational learning paradigms [4].

Shock electrodes were attached to the arm of each participant closest to the stimulus monitor, to maximize visibility of the arm to the other participant. Electrodes for recording electrodermal activity were attached to the distal phalanges of the middle and index fingers on the side not receiving shocks. The observer also received shock electrodes, as they would later be exposed to the threat of shocks during the testing phase, as is standard procedure in our observational learning protocol [4].

Each participant, in turn, went through an individual work-up procedure to calibrate the appropriate level of electrical stimulation in the experiment. Participants were instructed to choose a level of stimulation which they experienced as "uncomfortable, but not painful". Past research has shown that an observer receiving information about a (confederate) demonstrator's level of stimulation does not affect observational learning [6]. Both participants

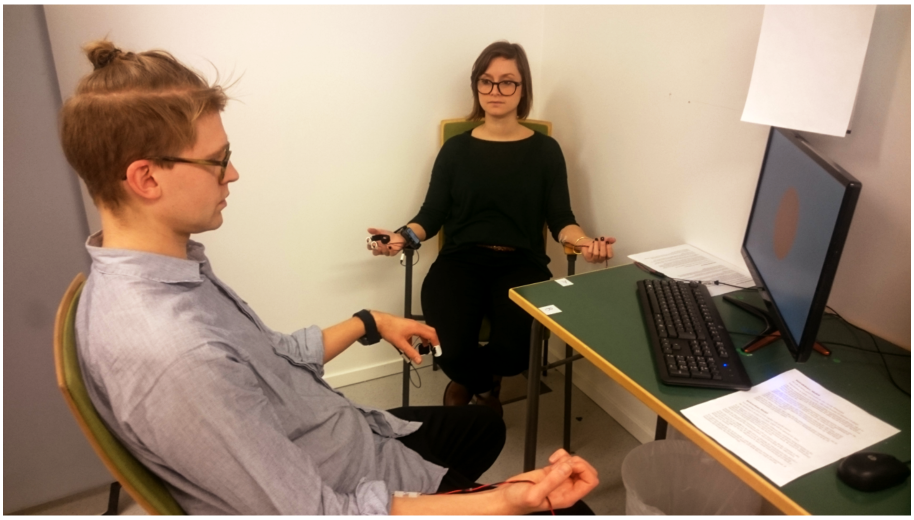

Figure 1: Picture of experimental setup during the learning phase. Observer (back) sits with a clear view of stimulus and the demonstrator. Demonstrator (front) fixates screen with stimulus presentation.

undergoing work-up implies that the observer underwent work-up prior to their first learning phase. This is a difference in our procedure to previous experiments and was included to facilitate the multiblock design of the present experiment (see below). Since the observers know the shock level they have set for themselves, they will have an accurate idea of the threat the shocks pose. In our previous, video-based work, observers had to guess what shocks would be like from the (exaggerated) reactions of the demonstrators. Hence, any effect of this prior shock exposure would likely have been to dampen the expression of the CS contingencies.

Once the participants were connected to the equipment the experiment commenced. The experimenter proceeded to give final verbal instructions to the participants. The demonstrator was reminded that the only way the observer could understand the pattern between the images that are shown on the computer screen and them receiving the shocks is by looking at them and therefore they are asked to act naturally and expressively, but without attempting to exaggerate or fake their responses. The demonstrator was additionally instructed to keep their gaze focused on the screen and to not look towards the observer. The observer was reminded that they may look wherever they wanted - to the demonstrator and the screen - to learn which stimulus predicts shocks and which doesn't. Finally, both participants were reminded that direct communication was forbidden. The experimenter remained in a corner of the room monitoring the participants, but also to ensure that participants were easily able to cancel their participation should they so wish.

Each block consisted of a learning phase and was followed immediately by a testing phase. In the learning phase the observer attempted to learn the CS contingencies by watching the demonstrators reactions to the CS images. The learning phase consisted of six alternating presentations of each CS+ and CS- image. Each CS was shown for six seconds. There was a variable length 10-16 second inter-trial interval between each CS presentation. Four of the six CS+ presentations, randomly determined, terminated with a shock to the demonstrator. Importantly, the observer received no shocks during this phase nor any instructions about which image was the CS+ (shock predicting) and which was the CS-. The only way to learn this contingency for the observer was through observation.

Following the learning phase, on-screen instructions informed the observer that they would view the same two CS images again and *now* receive shocks to the *same* image that they had observed the demonstrator previously receive shocks to [7, 8]. During this phase the demonstrator was instructed to close their eyes and a screen was placed between demonstrator and observer occluding the observers' view of the demonstrator (Supplementary Fig. 2). These steps were taken to ensure that the observer would not be able to pick up any cues during the testing phase about the valence of the CS images. Hence, any expression of heightened electrodermal activity to the CS+ image compared to the CS- image would *only* reflect associations observationally formed during the previous learning phase. During the testing phase each CS image was shown seven times. Unbeknownst to the observer, only the final CS+ presentation actually terminated with a shock. This final shock was given to ensure that the observers would consider the threat of shock credible also in the next block. The whole procedure was repeated the following block. Critically, each block featured *unique* stimulus images, hence there was no possibility of carry-over of learning between blocks.

After the second block had completed, the observer was asked to rate the demonstrator on four metrics: on how much pain the demonstrator seemed to be in, how much compassion the observer felt for the demonstrator, the extent to which the demonstrator was a good model and helped them learning and to how similar to them the demonstrator appeared to be. All ratings were completed using a 16cm visual-analog scale.

Once the observer had completed rating the demonstrator, the participants were asked to switch instruction sheets with one another. The experiment continued for two more blocks with the participants in reversed roles. Participants were unaware that this role reversal would occur. The role reversal was implemented to record more blocks from each recruited dyad. Ethical considerations limited us from exposing the demonstrator to more than two phases of direct conditioning. The role reversal means that each participant is observer for two of the four blocks of the experiment, and demonstrator for the other two. Half the participants begin the experiment

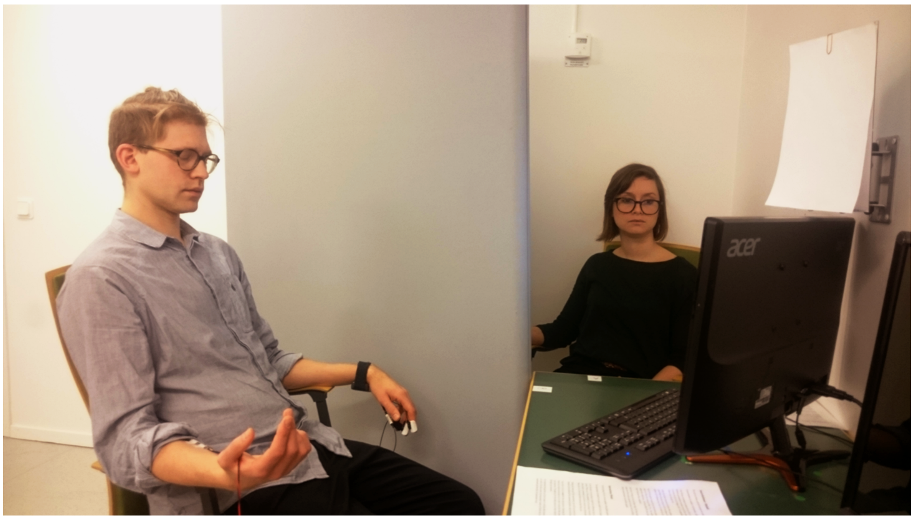

Figure 2: Picture of experimental setup during the testing phase. Observer (back) sits with a clear view of stimulus. Additional partition placed between observer and demonstrator occluding view. Demonstrator (front) has closed eyes and does not fixate stimulus screen.

with two blocks as observer while the other half of the participants become observers only after having been demonstrators first for two blocks.

Once all four blocks had completed, the current observer rated the demonstrator on the same four metrics as introduced above. Both participant then completed an Interpersonal Reactivity Index (IRI). The IRI consists of 28 items answered on a 5 point Likert scale ranging from "Does not describe me well" to "Describes me very well" and contains 4 subscales: Perspective taking, Fantasy, Empathic Concern and Personal Distress [9]. Finally, participants were thanked and debriefed.

#### Analysis

All analyses were performed in the R statistical language using the *brms* package [10]. We analyzed the data using Gaussian Bayesian multi-level regression models including varying intercepts and slopes by participant and between intercept and slope correlations. The outcome variable in all cases was the square-root transformed amplitude of each trial's skin conductance response, measured during the testing phase from the observer.

All categorical regressors were deviation coded (0.5/-0.5) and all continuous regressors were standardized. Weakly informative priors were used for all analyses to regularize estimates. These were Normal(0,0.5) for the intercept, Normal(0,0.2) for the slope of CS, and Normal(0,0.05) for all other slopes which usually contained the parameters of interest. LKJ(4) priors were placed on the correlation matrix and an Exponential(4) prior on the group standard deviations and and Exponential(2) prior on the model residual standard deviation.

Additionally, for two of the analyses we also fit hurdle lognormal models to the untransformed amplitude of the skin conductance responses. These are reported in Tables 2 and 7 below. Results from these analyses were consistent with the findings reported in the main paper.

For all estimates 95% credible intervals were computed as well as Bayes Factors based on the Savage-Dickey ratio for the parameter at the value 0. We rely on Bayes Factors to make inferences about effects. We interpret Bayes Factors above 10 to constitute strong evidence for an effect, and Bayes Factors between 3 and 10 to constitute weak evidence for an effect.

### Supplementary Results

#### Stability of synchrony across trials, blocks and participant roles

Analogously to our evaluation of overall threat learning, we tested if the predictive effect of synchrony on the observer's threat responses differed depending on if the observer started the experiment in that Role or, instead, as demonstrator and if it was the first or second Block in that role. It is possible that additional experience with the experiment might in some way affect how observers and demonstrators synchronize and that our effects were only present in parts of the data. Our analyses indicated that this was not the case. We found no evidence for moderation in the effect of synchrony on CS differentiation dependent on initial Role ($b$ = -0.007, $SE$ = 0.0017, $CrI$ = [-0.041, 0.025], $BF_{10}$ = 0.37) or by Block ($b$ = -0.004, $SE$ = 0.015, $CrI$ = [-0.033, 0.024], $BF_{10}$ = 0.31). In other words, the relationship between synchrony and strength of conditioned responses was not affected by participants switching roles or by their experience with the observational learning procedure. See also Table 8 below.

We next tested if the effect of synchrony (PC1) changed over the series of consecutive CS image presentations as the participants' responses extinguished (cf. Fig. 1) . We added a Trial variable to a model with the CS status and the synchrony component predictors, together with all their interactions. Unsurprisingly there was both a main effect of trial ($b$ = -0.034, $SE$ = 0.003, $CrI$ = [-0.040, -0.029], $BF_{10} > 10^6$) and a weak interaction with CS status possibly indicating marginally faster extinction of responses to the CS+ ($b$ = -0.013, $SE$ = 0.005, $CrI$ = [-0.023, -0.003], $BF_{10}$ = 3.0). However, of critical interest, the relationship between synchrony and CS differentiation did not interact with trial, with strong evidence for null effect ($b$ = 0.0015, $SE$ = 0.003, $CrI$ = [-0.005, 0.008], $BF_{10}$ = 0.070). The relationship between CS differentiation and synchrony was stable across all trials in the testing phase. See Table 9 below.

Table 1: Results investigating CS differentiation and potential moderators of Role and Block.

| | Estimate | Est.Error | Q2.5 | Q97.5 | $BF_{01}$ | $BF_{10}$ |
|---|---|---|---|---|---|---|
| intercept | 0.2845 | 0.0154 | 0.2538 | 0.3145 | | |
| Role | -0.0124 | 0.0260 | -0.0633 | 0.0380 | 1.7125 | 0.5839 |
| CS | 0.1533 | 0.0165 | 0.1203 | 0.1855 | -0.0000 | $> 10^6$ |
| Block | -0.0003 | 0.0109 | -0.0217 | 0.0211 | 4.5753 | 0.2186 |
| Role:CS | 0.0356 | 0.0276 | -0.0182 | 0.0903 | 0.7958 | 1.2566 |
| Role:Block | -0.0182 | 0.0202 | -0.0580 | 0.0221 | 1.5866 | 0.6303 |
| CS:Block | 0.0303 | 0.0187 | -0.0066 | 0.0668 | 0.7452 | 1.3419 |
| Role:CS:Block | 0.0011 | 0.0318 | -0.0624 | 0.0636 | 1.6389 | 0.6102 |

Table 2: Results from hurdle lognormal model investigating CS differentiation and potential moderators of Role and Block. Coefficients from main model on log scale. Role, CS status and Block modeled in hurdle component, prefixed *hu*. Hurdle coefficients on logit scale.

| | Estimate | Est.Error | Q2.5 | Q97.5 | $BF_{01}$ | $BF_{10}$ |
|---|---|---|---|---|---|---|
| intercept | -3.1801 | 0.1079 | -3.3946 | -2.9709 | | |
| Role | -0.1145 | 0.1979 | -0.5037 | 0.2743 | 2.0919 | 0.4780 |
| CS | 0.8993 | 0.1008 | 0.7016 | 1.0986 | 0.0000 | $> 10^6$ |
| Block | -0.2047 | 0.0814 | -0.3629 | -0.0441 | 0.2688 | 3.7203 |
| Role:CS | 0.1442 | 0.1887 | -0.2281 | 0.5160 | 1.9931 | 0.5017 |
| Role:Block | -0.2464 | 0.1551 | -0.5535 | 0.0568 | 0.9024 | 1.1081 |
| CS:Block | 0.2503 | 0.1436 | -0.0321 | 0.5284 | 0.7612 | 1.3136 |
| Role:CS:Block | -0.0920 | 0.2539 | -0.5935 | 0.4111 | 1.8327 | 0.5456 |
| hu intercept | -1.5143 | 0.0990 | -1.7135 | -1.3219 | | |
| hu Role | 0.0300 | 0.1375 | -0.2383 | 0.3010 | 1.4867 | 0.6726 |
| hu CS | -0.8008 | 0.1131 | -1.0264 | -0.5842 | -0.0000 | $> 10^6$ |
| hu Block | -0.0120 | 0.0907 | -0.1867 | 0.1677 | 2.2527 | 0.4439 |

Table 3: Results from *separate* regression models, each investigating the DETerminism, LAMinarity, maximum diagonal line Length and relative ENTRopy CRQA components, respectively.

| | Estimate | Est.Error | Q2.5 | Q97.5 | $BF_{01}$ | $BF_{10}$ |
|---|---|---|---|---|---|---|
| intercept | 0.2858 | 0.0156 | 0.2550 | 0.3157 | | |
| CS | 0.1537 | 0.0160 | 0.1227 | 0.1853 | 0.0000 | $> 10^6$ |
| DET | 0.0085 | 0.0089 | -0.0087 | 0.0259 | 3.7284 | 0.2682 |
| CS:DET | 0.0553 | 0.0134 | 0.0294 | 0.0818 | 0.0016 | 610.0279 |
| intercept | 0.2851 | 0.0156 | 0.2546 | 0.3159 | | |
| CS | 0.1522 | 0.0160 | 0.1206 | 0.1834 | 0.0000 | $> 10^6$ |
| LAM | 0.0075 | 0.0080 | -0.0083 | 0.0232 | 3.9600 | 0.2525 |
| CS:LAM | 0.0511 | 0.0136 | 0.0245 | 0.0777 | 0.0007 | 1481.2072 |
| intercept | 0.2851 | 0.0153 | 0.2549 | 0.3152 | | |
| CS | 0.1540 | 0.0163 | 0.1217 | 0.1857 | 0.0000 | $> 10^6$ |
| maxL | 0.0009 | 0.0075 | -0.0138 | 0.0155 | 6.6064 | 0.1514 |
| CS:maxL | 0.0368 | 0.0136 | 0.0100 | 0.0634 | 0.1031 | 9.6968 |
| intercept | 0.2849 | 0.0154 | 0.2546 | 0.3153 | | |
| CS | 0.1531 | 0.0170 | 0.1194 | 0.1865 | -0.0000 | $> 10^6$ |
| rENTR | -0.0014 | 0.0068 | -0.0148 | 0.0118 | 7.3602 | 0.1359 |
| CS:rENTR | 0.0268 | 0.0119 | 0.0032 | 0.0500 | 0.3637 | 2.7498 |

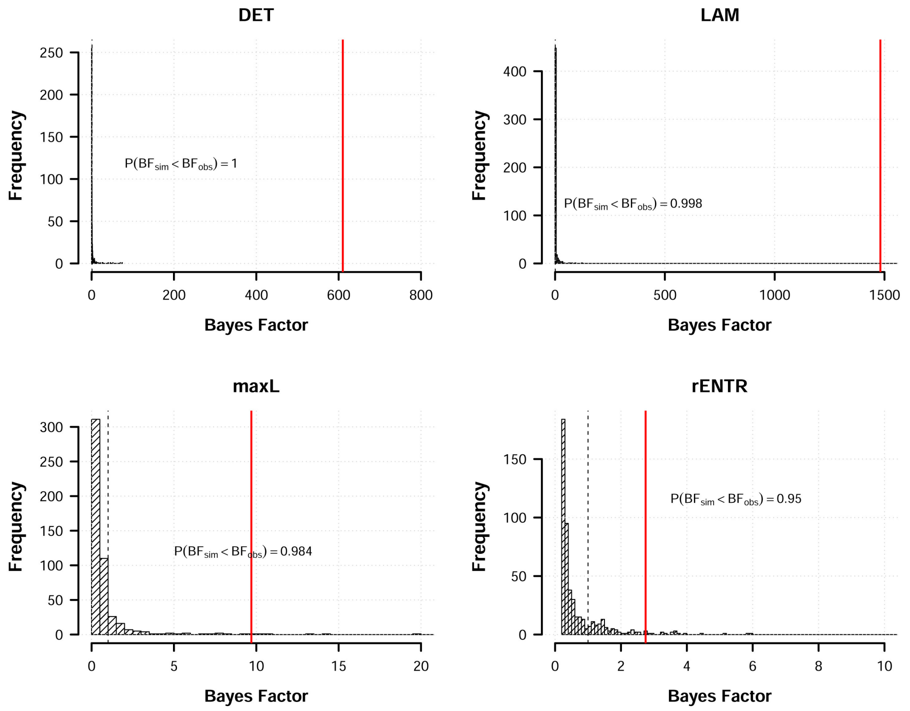


Figure 3: Histograms of model evidence (Bayes Factors) in favor of an effect of each of the four CRQA metrics on CS differentiation, computed from creating pseudo-dyads through permutation of the real participants pairings in our dataset. The vertical red line indicates the observed Bayes Factor in our actual dyads and the dashed vertical line shows BF = 1.

Table 4: Results from model *jointly* regressing the DETerminism, LAMinarity, maximum diagonal line Length and relative ENTRopy CRQA components.

| | Estimate | Est.Error | Q2.5 | Q97.5 | $BF_{01}$ | $BF_{10}$ |
|---|---|---|---|---|---|---|
| intercept | 0.2856 | 0.0155 | 0.2547 | 0.3157 | | |
| CS | 0.1534 | 0.0164 | 0.1209 | 0.1857 | 0.0000 | $> 10^6$ |
| DET | 0.0106 | 0.0150 | -0.0187 | 0.0399 | 2.6145 | 0.3825 |
| maxL | -0.0066 | 0.0104 | -0.0269 | 0.0137 | 4.0370 | 0.2477 |
| rENTR | -0.0079 | 0.0092 | -0.0260 | 0.0102 | 3.8983 | 0.2565 |
| LAM | 0.0085 | 0.0120 | -0.0151 | 0.0321 | 3.1064 | 0.3219 |
| CS:DET | 0.0315 | 0.0214 | -0.0107 | 0.0732 | 0.7844 | 1.2748 |
| CS:maxL | 0.0092 | 0.0165 | -0.0237 | 0.0410 | 2.7304 | 0.3662 |
| CS:rENTR | 0.0050 | 0.0150 | -0.0242 | 0.0350 | 3.2494 | 0.3077 |
| CS:LAM | 0.0262 | 0.0177 | -0.0082 | 0.0612 | 0.8907 | 1.1228 |

Table 5: Factor loadings of each principal component extracted from the CRQA metric as well as their explained variance.

| CRQA Metrics | PC1 | PC2 | PC3 | PC4 |
|---|---|---|---|---|
| DET | 0.586 | -0.101 | 0.111 | 0.796 |
| maxL | 0.444 | -0.697 | 0.323 | -0.46 |
| rENTR | 0.428 | 0.697 | 0.494 | -0.295 |
| LAM | 0.525 | 0.135 | -0.799 | -0.258 |
| Variance Explained | 0.631 | 0.214 | 0.108 | 0.047 |

Table 6: Results from regression model using principal components of CRQA metrics.

| | Estimate | Est.Error | Q2.5 | Q97.5 | $BF_{01}$ | $BF_{10}$ |
|---|---|---|---|---|---|---|
| intercept | 0.2844 | 0.0153 | 0.2540 | 0.3146 | | |
| CS | 0.1554 | 0.0160 | 0.1242 | 0.1867 | 0.0000 | $> 10^6$ |
| PC1 | 0.0030 | 0.0054 | -0.0077 | 0.0137 | 8.1161 | 0.1232 |
| PC2 | -0.0018 | 0.0075 | -0.0163 | 0.0130 | 6.5536 | 0.1526 |
| PC3 | -0.0120 | 0.0127 | -0.0366 | 0.0132 | 2.4133 | 0.4144 |
| PC4 | 0.0156 | 0.0193 | -0.0227 | 0.0537 | 1.8955 | 0.5276 |
| CS:PC1 | 0.0381 | 0.0092 | 0.0205 | 0.0564 | 0.0009 | 1156.4511 |
| CS:PC2 | -0.0029 | 0.0128 | -0.0281 | 0.0222 | 3.9450 | 0.2535 |
| CS:PC3 | -0.0128 | 0.0200 | -0.0522 | 0.0270 | 2.0739 | 0.4822 |
| CS:PC4 | 0.0130 | 0.0263 | -0.0389 | 0.0642 | 1.6544 | 0.6044 |

Table 7: Results from regression hurdle lognormal model using principal components of CRQA metrics. Coefficients from lognormal component on log scale. CS status modeled in hurdle component, prefixed *hu*. Hurdle coefficients on logit scale.

| | Estimate | Est.Error | Q2.5 | Q97.5 | $BF_{01}$ | $BF_{10}$ |
|---|---|---|---|---|---|---|
| intercept | -3.1714 | 0.1084 | -3.3828 | -2.9569 | | |
| CS | 0.8941 | 0.0962 | 0.7086 | 1.0868 | 0.0000 | $> 10^6$ |
| PC1 | -0.0164 | 0.0469 | -0.1105 | 0.0743 | 10.3555 | 0.0966 |
| PC2 | -0.0445 | 0.0631 | -0.1694 | 0.0787 | 6.1885 | 0.1616 |
| PC3 | -0.1622 | 0.1041 | -0.3661 | 0.0432 | 1.5067 | 0.6637 |
| PC4 | -0.0543 | 0.1578 | -0.3640 | 0.2571 | 3.0311 | 0.3299 |
| CS:PC1 | 0.2417 | 0.0593 | 0.1252 | 0.3575 | 0.0000 | 71647.2706 |
| CS:PC2 | 0.0807 | 0.1050 | -0.1213 | 0.2882 | 3.6482 | 0.2741 |
| CS:PC3 | -0.0995 | 0.1335 | -0.3630 | 0.1592 | 2.8221 | 0.3543 |
| CS:PC4 | 0.0224 | 0.2006 | -0.3788 | 0.4168 | 2.5501 | 0.3921 |
| hu intercept | -1.4928 | 0.0966 | -1.6884 | -1.3102 | | |
| hu CS | -0.7951 | 0.1116 | -1.0213 | -0.5792 | 0.0000 | $> 10^6$ |

Table 8: Results from model investigating if Role and Block moderate first synchrony component (PC1) effect on CS differentiation.

| | Estimate | Est.Error | Q2.5 | Q97.5 | $BF_{01}$ | $BF_{10}$ |
|---|---|---|---|---|---|---|
| intercept | 0.2823 | 0.0154 | 0.2522 | 0.3127 | | |
| CS | 0.1548 | 0.0164 | 0.1227 | 0.1871 | 0.0000 | $> 10^6$ |
| Role | -0.0115 | 0.0255 | -0.0610 | 0.0383 | 1.7509 | 0.5711 |
| PC1 | 0.0027 | 0.0060 | -0.0091 | 0.0144 | 7.7750 | 0.1286 |
| Block | -0.0025 | 0.0112 | -0.0244 | 0.0196 | 4.4405 | 0.2252 |
| CS:PC1 | 0.0353 | 0.0093 | 0.0170 | 0.0540 | 0.0028 | 356.6022 |
| CS:Role | 0.0239 | 0.0270 | -0.0288 | 0.0774 | 1.3167 | 0.7594 |
| CS:Block | 0.0242 | 0.0188 | -0.0125 | 0.0611 | 1.1572 | 0.8641 |
| Role:Block | -0.0272 | 0.0209 | -0.0682 | 0.0142 | 0.6455 | 1.5492 |
| Role:PC1 | 0.0226 | 0.0116 | -0.0004 | 0.0453 | 4.2453 | 0.2356 |
| PC1:Block | 0.0067 | 0.0088 | -0.0105 | 0.0241 | 0.9989 | 1.0011 |
| CS:Role:PC1 | -0.0069 | 0.0168 | -0.0405 | 0.0253 | 2.7339 | 0.3658 |
| CS:PC1:Block | -0.0042 | 0.0146 | -0.0327 | 0.0244 | 3.2075 | 0.3118 |

Table 9: Results from model investigating if first synchrony component (PC1) effect is moderated by consecutive CS presentations (Trial)

| | Estimate | Est.Error | Q2.5 | Q97.5 | $BF_{01}$ | $BF_{10}$ |
|---|---|---|---|---|---|---|
| intercept | 0.3875 | 0.0197 | 0.3485 | 0.4257 | | |
| CS | 0.1930 | 0.0230 | 0.1485 | 0.2387 | 0.0000 | $> 10^6$ |
| PC1 | 0.0047 | 0.0073 | -0.0096 | 0.0191 | 5.5919 | 0.1788 |
| Trial | -0.0343 | 0.0029 | -0.0400 | -0.0285 | -0.0000 | $> 10^6$ |
| CS:PC1 | 0.0347 | 0.0125 | 0.0103 | 0.0596 | 0.0783 | 12.7677 |
| CS:Trial | -0.0131 | 0.0050 | -0.0229 | -0.0033 | 0.3333 | 3.0002 |
| PC1:Trial | -0.0003 | 0.0017 | -0.0037 | 0.0031 | 27.5662 | 0.0363 |
| CS:PC1:Trial | 0.0015 | 0.0030 | -0.0045 | 0.0075 | 14.2071 | 0.0704 |

Table 10: Results investigating specificity of synchrony component predicting CS differentiation using three alternative predictors. Social UCS: average strength of the observer's skin conductance response to the demonstrator receiving shocks. CS learn: average difference of the observer's responses to the CS+ over CS- during the learning phase. Cor: time-lagged correlation between demonstrator's and observer's skin conductance time-series during the learning phase.

| | Estimate | Est.Error | Q2.5 | Q97.5 | $BF_{01}$ | $BF_{10}$ |
|---|---|---|---|---|---|---|
| intercept | 0.2898 | 0.0142 | 0.2621 | 0.3179 | | |
| CS | 0.1581 | 0.0165 | 0.1258 | 0.1899 | 0.0000 | $> 10^6$ |
| social UCS | 0.0450 | 0.0123 | 0.0207 | 0.0691 | 0.0065 | 154.3204 |
| CS learn | -0.0088 | 0.0104 | -0.0287 | 0.0125 | 3.0758 | 0.3251 |
| Cor | -0.0060 | 0.0090 | -0.0238 | 0.0118 | 4.5578 | 0.2194 |
| PC1 | 0.0005 | 0.0067 | -0.0127 | 0.0135 | 7.7961 | 0.1283 |
| CS:social UCS | 0.0211 | 0.0171 | -0.0123 | 0.0542 | 1.3847 | 0.7222 |
| CS:CS learn | -0.0138 | 0.0162 | -0.0446 | 0.0192 | 2.1424 | 0.4668 |
| CS:Cor | 0.0072 | 0.0135 | -0.0194 | 0.0336 | 3.2450 | 0.3082 |
| CS:PC1 | 0.0339 | 0.0100 | 0.0146 | 0.0538 | 0.0140 | 71.4138 |
| social UCS:PC1 | -0.0059 | 0.0087 | -0.0229 | 0.0109 | 4.6498 | 0.2151 |
| CS learn:PC1 | 0.0023 | 0.0084 | -0.0137 | 0.0194 | 5.8423 | 0.1712 |
| Cor:PC1 | -0.0050 | 0.0063 | -0.0174 | 0.0073 | 6.0158 | 0.1662 |
| CS:social UCS:PC1 | -0.0002 | 0.0122 | -0.0243 | 0.0238 | 4.1555 | 0.2406 |
| CS:CS learn:PC1 | 0.0106 | 0.0125 | -0.0135 | 0.0353 | 2.8321 | 0.3531 |
| CS:Cor:PC1 | -0.0051 | 0.0100 | -0.0249 | 0.0147 | 4.3974 | 0.2274 |

Table 11: Means and standard deviations of participants' scores on the four sub-scales of the interpersonal reactivity index and their ratings of the Demonstrator.

| Sub-scale | Mean | Std. Dev. |
|---|---|---|
| Personal Distress | 12.4 | 4.6 |
| Perspective Taking | 18.1 | 4.6 |
| Empathic Concerns | 19.8 | 4.1 |
| Fantasy | 18.5 | 5.4 |
| Rating | Mean | Std. Dev. |
| Pain | 7.1 | 3.2 |
| Compassion | 7.9 | 3.9 |
| Good Model | 11.4 | 3.7 |
| Similarity | 8.8 | 3.9 |

Table 12: Results investigating effects of self-rated empathy on CS differentiation and moderation of synchrony component (PC1) on CS differentiation. Four sub-scales from interpersonal reactivity index (IRI), FS: Fantasy; EC: Empathic concern; PT: Perspective taking; PD: Personal Distress.

| | Estimate | Est.Error | Q2.5 | Q97.5 | $BF_{01}$ | $BF_{10}$ |
|---|---|---|---|---|---|---|
| intercept | 0.2794 | 0.0148 | 0.2508 | 0.3089 | | |
| CS | 0.1506 | 0.0162 | 0.1188 | 0.1820 | 0.0000 | $> 10^6$ |
| IRI_FS | 0.0043 | 0.0159 | -0.0268 | 0.0355 | 3.0395 | 0.3290 |
| IRI_EC | 0.0009 | 0.0172 | -0.0334 | 0.0344 | 2.9652 | 0.3372 |
| IRI_PT | -0.0232 | 0.0163 | -0.0549 | 0.0089 | 1.0767 | 0.9287 |
| IRI_PD | -0.0331 | 0.0154 | -0.0633 | -0.0032 | 0.3081 | 3.2452 |
| PC1 | 0.0042 | 0.0062 | -0.0080 | 0.0164 | 6.3536 | 0.1574 |
| CS:PC1 | 0.0322 | 0.0093 | 0.0142 | 0.0504 | 0.0095 | 104.7668 |
| CS:IRI_FS | 0.0000 | 0.0175 | -0.0341 | 0.0341 | 2.8879 | 0.3463 |
| CS:IRI_EC | -0.0043 | 0.0174 | -0.0382 | 0.0293 | 2.7199 | 0.3677 |
| CS:IRI_PT | -0.0112 | 0.0182 | -0.0471 | 0.0245 | 2.2823 | 0.4382 |
| CS:IRI_PD | -0.0071 | 0.0172 | -0.0408 | 0.0269 | 2.6982 | 0.3706 |
| IRI_FS:PC1 | -0.0062 | 0.0076 | -0.0212 | 0.0086 | 4.9436 | 0.2023 |
| IRI_EC:PC1 | 0.0077 | 0.0075 | -0.0074 | 0.0221 | 3.7156 | 0.2691 |
| IRI_PT:PC1 | 0.0070 | 0.0079 | -0.0084 | 0.0229 | 4.5541 | 0.2196 |
| IRI_PD:PC1 | -0.0056 | 0.0069 | -0.0192 | 0.0080 | 5.1503 | 0.1942 |
| CS:IRI_FS:PC1 | -0.0048 | 0.0110 | -0.0264 | 0.0170 | 4.2484 | 0.2354 |
| CS:IRI_EC:PC1 | 0.0070 | 0.0101 | -0.0129 | 0.0268 | 3.8578 | 0.2592 |
| CS:IRI_PT:PC1 | -0.0030 | 0.0110 | -0.0251 | 0.0184 | 4.2944 | 0.2329 |
| CS:IRI_PD:PC1 | -0.0071 | 0.0101 | -0.0268 | 0.0128 | 3.9362 | 0.2541 |

Table 13: Results investigating effects of observer's rating of demonstrator on CS differentiation and moderation of synchrony component (PC1) on CS differentiation.

| | Estimate | Est.Error | Q2.5 | Q97.5 | $BF_{01}$ | $BF_{10}$ |
|---|---|---|---|---|---|---|
| intercept | 0.2898 | 0.0157 | 0.2591 | 0.3204 | | |
| CS | 0.1606 | 0.0165 | 0.1279 | 0.1931 | -0.0000 | $> 10^6$ |
| Sim | -0.0159 | 0.0169 | -0.0494 | 0.0171 | 1.9889 | 0.5028 |
| Pain | -0.0187 | 0.0172 | -0.0522 | 0.0158 | 1.5624 | 0.6400 |
| Comp | -0.0084 | 0.0176 | -0.0434 | 0.0263 | 2.4367 | 0.4104 |
| GoodModel | 0.0184 | 0.0186 | -0.0184 | 0.0548 | 1.6623 | 0.6016 |
| PC1 | 0.0032 | 0.0062 | -0.0089 | 0.0154 | 6.8756 | 0.1454 |
| CS:PC1 | 0.0359 | 0.0095 | 0.0174 | 0.0548 | 0.0069 | 145.3607 |
| CS:Sim | 0.0293 | 0.0178 | -0.0057 | 0.0642 | 0.6966 | 1.4356 |
| CS:Pain | 0.0054 | 0.0181 | -0.0302 | 0.0403 | 2.6990 | 0.3705 |
| CS:Comp | -0.0075 | 0.0182 | -0.0431 | 0.0286 | 2.5716 | 0.3889 |
| CS:GoodModel | 0.0046 | 0.0189 | -0.0325 | 0.0415 | 2.5510 | 0.3920 |
| Sim:PC1 | -0.0049 | 0.0073 | -0.0192 | 0.0097 | 5.4679 | 0.1829 |
| Pain:PC1 | -0.0036 | 0.0066 | -0.0166 | 0.0096 | 6.5705 | 0.1522 |
| Comp:PC1 | 0.0061 | 0.0077 | -0.0094 | 0.0209 | 4.6628 | 0.2145 |
| GoodModel:PC1 | 0.0034 | 0.0078 | -0.0116 | 0.0192 | 6.0963 | 0.1640 |
| CS:Sim:PC1 | -0.0048 | 0.0113 | -0.0271 | 0.0174 | 4.1712 | 0.2397 |
| CS:Pain:PC1 | 0.0036 | 0.0105 | -0.0172 | 0.0242 | 4.4863 | 0.2229 |
| CS:Comp:PC1 | -0.0140 | 0.0120 | -0.0379 | 0.0088 | 2.1178 | 0.4722 |
| CS:GoodModel:PC1 | 0.0046 | 0.0119 | -0.0182 | 0.0287 | 4.0661 | 0.2459 |

[3] Wolfram Boucsein. *Electrodermal activity*. Springer Science & Business Media, 2012.
